# Cortical surface electrical potentials are composed of multiple bandlimited frequency components, including high-gamma

**DOI:** 10.1101/2022.07.22.501176

**Authors:** Jesse A. Livezey, Ahyeon Hwang, Kseniya Usovich, Maximilian E. Dougherty, Edward F. Chang, Kristofer E. Bouchard

## Abstract

A common challenge in neuroscience is how to decompose noisy, multi-source signals measured in experiments into biophysically interpretable components. Analysis of cortical surface electrical potentials (CSEPs) measured using electrocorticography arrays (ECoG) typifies this problem. We hypothesized that high frequency (70-1,000 Hz) CSEPs are composed of broadband (i.e., power-law) and bandlimited components with potentially differing biophysical origins. In particular, the high-gamma band (70-150 Hz) has been shown to be highly predictive for encoding and decoding behaviors and stimuli. Despite its demonstrated importance, whether high-gamma is composed of a bandlimited signal is poorly understood. To address this gap, we recorded CSEPs from rat auditory cortex and demonstrate that the evoked CSEPs are composed of multiple distinct frequency components, including high-gamma. We then show, using a novel robust regression method, that at fast timescales and on single trials during speech production, human high-gamma amplitude cannot be explained by a modulating power-law component; thus, high-gamma is band-limited. Furthermore, we show that the power-law component is less predictive of produced speech compared to the raw high-gamma amplitude. Finally, we show that the largest variance component of human ECoG signals is low-frequency and band-limited, not broadband. Together these results demonstrate that there are multiple, band-limited components of high frequency power in cortical surface electrical potentials, including the high-gamma band, which may have different biophysical origins.

**Significance Statement:** Electrocortigraphy (ECoG) records cortical surface electrical potentials (CSEPs). ECoG is utilized in both humans and animal models to understand distributed cortical processing and for brain machine interfaces. The spectral structure of evoked CSEPs is greatly debated. To address this issue, we recorded from rat auditory cortex using *µ*ECoG and human ventral sensory-motor cortex during speech production with high-density ECoG. Using novel analytic approaches, we found that evoked CSEPs are composed of multiple band-limited components, including high-gamma. These results contrast with the dominant thinking in the field that the high-frequency power in ECoG is broadband. Our results raise the possibility that distinct frequency components of ECoG are biomarkers of processing in different cortical layers.

## Introduction

Understanding the spectral structure of time-varying neural recordings, such as cortical surface electrical potentials (CSEPs) recorded using ECoG, is important for linking the biophysical sources of electrical potentials to perception, behavior, and disease in humans and animal models. ECoG records cortical surface electrical potentials (CSEPs), which, like all electrical recordings in the brain, reflect a weighted superposition of all electrical sources surrounding the electrode (Buzsáki, Anastassiou, et al. 2012; Einevoll et al. 2013). Because of the 1*/f*^*m*^ fall-off of power with frequency of brain signals (Buzsáki, Anastassiou, et al. 2012; Einevoll et al. 2013; Freeman and Zhai 2009; Miller, Sorensen, et al. 2009), many studies of ECoG and local field potentials (LFP) focus on lower frequencies (e.g., *<*60Hz). The high frequency range of CSEPs (70-1,000 Hz), and in particular the high-gamma band (70-150Hz), is of considerable interest due to its correlation with spiking activity (Ray and Maunsell 2011), its columnar localization and laminar origin (Baratham et al. 2022), and its potential use as a highly predictive clinical signal for human brain-computer interfaces (Bouchard and Chang 2014b; Livezey, Bouchard, et al. 2019; Livezey and Glaser 2021; Moses et al. 2019; Mugler et al. 2014). A central, unresolved question is whether the high frequency range of CSEPs, including high-gamma, is a single broadband component or is composed of a combination of broadband and band-limited components.

The amplitude of the high-gamma band, and high frequency amplitudes more broadly, are commonly thought to reflect a single broadband component in the CSEP, potentially with a 1*/f*^*m*^ form (Manning et al. 2009; Miller, Honey, et al. 2014; Miller, Leuthardt, et al. 2007; Miller, Sorensen, et al. 2009; Miller, Zanos, et al. 2009). In contrast, a lone study has reported differential encoding of speech tasks by high-frequencies in humans (Gaona et al. 2011). Previous work has shown that the high-gamma amplitude is separate from lower frequency oscillations including the gamma oscillation (Crone, Korzeniewska, et al. 2011; Crone, Miglioretti, et al. 1998; Ray and Maunsell 2011). High-gamma amplitude is differentially correlated to lower frequency oscillations depending on behavioral state and the functional area of cortex (Alekseichuk et al. 2016; Canolty et al. 2010; Livezey, Bouchard, et al. 2019). Importantly for clinical brain-computer interface applications, it has been shown that the CSEP high-gamma amplitude is a highly predictive signal for decoding movements, as well as spoken and perceived speech (Anumanchipalli et al. 2019; Bouchard and Chang 2014b; Livezey, Bouchard, et al. 2019; Mesgarani et al. 2014; Mugler et al. 2014; Pei et al. 2011). However, it remains unresolved whether the fast timescale modulations in high frequency CSEPs, including the high-gamma amplitude, correspond to a single time-varying broadband component or a combination of a broadband and one-or-more band-limited components. Importantly, it is the fast time-scale (10’s of milliseconds) modulations of high-frequency amplitudes that contains task relevant information. Distinguishing whether high frequencies in CSEPS are band-limited or broad-band impacts their biophysical interpretation, for example whether the recorded signals are differentially generated (and potentially resolvable) across layers of cortex (Baratham et al. 2022).

Several technical issues have impeded a definitive answer to this question. In order to estimate the broadband component in CSEPS, previous studies have used basic 1/f power-law regression (Donoghue et al. 2020; Manning et al. 2009; Miller, Sorensen, et al. 2009) and a non-standard version of PCA (Miller, Honey, et al. 2014; Miller, Zanos, et al. 2009) to extract the broadband component. Furthermore, the spectral structure of high-frequency CSEPs has mainly been studied over long time windows, which may average out temporally-sparse (i.e., task selective) high frequency activity, and also limits the study of behavioral-timescale modulations. CSEPs recorded using ECoG arrays are most commonly recorded in humans using clinical recording hardware which often has a noise floor which interferes with measuring signals above 200 Hz (Miller, Sorensen, et al. 2009). Finally, clinical electrodes are commonly relatively large compared to a cortical column which may average out functionally-sparse high frequency activity. Due to these difficulties, the spectral structure of time-varying, high frequency CSEPs is poorly understood.

Based on observations from our recently published study (Baratham et al. 2022), we hypothesized that high frequency CSEPs are composed of one-or-more bandlimited components in addition to a broadband component. To test this hypothesis, we used high-quality neural amplifiers to record with *µ*ECoG from rat auditory cortex in response to sounds and high-density ECoG from human sensorimotor cortex during speech production. We developed and deployed advanced analytic techniques, which allowed us to demonstrate high frequency CSEPs are composed of multiple distinct, time-varying components.

## Materials and Methods

### Rat auditory experiments

Frequencies beyond 200 Hz are often difficult to resolve in human ECoG recordings due to limitations of clinical amplifiers or the relatively large size of electrodes. In order to gain access to the high frequency range (70-1,000 Hz) of cortical surface electrical potentials (CSEPs), we recorded CSEPs from rat auditory cortex using *µ*ECoG arrays. Data collection and preprocessing methodology for the rat data has been previously described (Baratham et al. 2022; Dougherty et al. 2019). All animal procedures were performed in accordance with established animal care protocols approved by the Lawrence Berkeley National Laboratory Institutional Animal Care and Use Committee.

Six female Sprague Dawley rats were used in this study. We performed experiments in six anesthetized female Sprague Dawley rats. Animals were given a 1 mg/kg subcutaneous (s.q.) injection of Dexamethasone the night before a procedure to reduce cerebral edema. An anesthetic state was established with an inductive dose of ketamine (95 mg/kg i.p.) and xylazine (10 mg/kg i.p.). Anesthetic state was assessed using toe pinch reflex and monitoring respiration rate. Additional doses of ketamine (55 mg/kg i.p.) and xylazine (5 mg/kg i.p.) were administered as needed to maintain a negative reflex and a regular respiration rate. Respiration was supported with a preoperative subcutaneous injection of atropine (0.2 mg/kg for rats), and a perioperative nose cone supplying 0.8 L/min of O_2_. A water heating bed provided thermostatic regulation. To prevent dehydration over the 10-hour surgery and recording session, subcutaneous saline injections (1 ml/kg) were provided every 3 hours. Once anesthetized the rodent was affixed to a snout stereotax without earbars. After stable anesthetic state was achieved, an incision was made along the sagittal midline. All the soft tissue on top of the skull was removed to reveal the lambda and bregma fissures. Two 1mm burr holes were drilled over non-auditory cortical areas–one between the lambda and bregma on the left hemisphere, and another anterior to bregma on the right hemisphere. These serve to reduce intracranial pressure and provide a reference for electrophysiological recordings. The right masseter muscle was then transected to uncover the portion of cranium lying over the right auditory cortex. Using a 1 mm diamond tapered round Stryker dental drill, a craniotomy was performed to expose the cortex.

All neural data were recorded with a multi-channel amplifier optically connected to a digital signal processor (Tucker-Davis Technologies, TDT, Alachua, FL). Rodent data was acquired at 12 kHz and lowpass filtered to the Nyquist frequency (6 kHz). Rodent electrophysiological recordings were made with custom designed 64 or 128-channel *µ*ECoG grids. These arrays were placed over the primary auditory cortex as identified by anatomical and physiological properties. The dura was surgically removed and the *µ*ECoG grid was placed directly on the pial surface. The *µ*ECoG grids were placed over the primary auditory cortex and grounded via a silver wire inserted into a non-auditory cortical area in the contra-lateral hemisphere, which also served as the reference. Each contact on the grid had an impedance of 30 ± 10 kΩ after electroplating with platinum black (measured in 1x phosphate-buffered saline at 1 kHz), and had an exposed diameter of 40 *µ*m, with a 200 *µ*m inter-electrode pitch.

### White noise auditory Stimuli

White noise bursts (*N* = 60 bursts, 100 ms, frequency content from [1-50 kHz]) were played back once per second through a DVD player (Sony), attached to a TDT amplifier and played through an electrostatic speaker located near the left ear of the animal (80 dB SPL, A-scale).

### Site selection for rat *µ*ECoG

We recorded from the surface of auditory cortex at 17 *µ*ECoG grid placements across 6 rats in response to the white noise stimuli. This yielded 1664 total recording sites. Of these, *N* = 1402 (84%) were deemed to have an auditory response (peak z-scored response in any neural frequency bin *>* 3.0). These responsive sites were subject to further analysis of cortical surface electrical potentials (CSEPs). Of these sites, *N* = 1156 (83% of responsive sites, 70% of total sites) were deemed to have an auditory evoked multi-unit evoked response (z-scored of tMuA response *>* 0.5).

### Spectral analysis of rat cortical surface electrical potentials (CSEPs)

We calculated the spectrogram of the entire recorded electrical potential time series for each electrode from 4-1200 Hz (54 bins) using a Matlab package for constant-Q wavelet transform (Schörkhuber and Klapuri 2010). Constant-Q refers to a time-frequency decomposition in which frequency bins are geometrically spaced and Q-factors (ratios of the center frequencies to bandwidths) are equal for all bins. The non-causal component of displayed responses is due to the large bandwidth at lower frequencies of the constant-Q time-frequency transform. The amplitude for each frequency bin in the neural signal was normalized relative to baseline statistics by z-scoring. The baseline statistics were computed from the periods of silence between each stimulus presentation. Z-scoring largely removes the canonical 1/f fall off of power with frequency (characteristic of many natural signals), highlighting stimulus evoked changes (**Fig. 2**). Responses to stimuli were taken as the average z-scored activity ±5 ms around the peak response time after the onset of the auditory stimulus. The shielding and grounding of our rodent experimental recording systems was sufficient to avoid any significant 60 Hz noise in the *µ*ECoG recordings. We determined z-scores for the canonical neural frequency bands by taking the mean z-scores across frequency bins in the corresponding frequency range: Beta (10-27 Hz), Gamma (30-57 Hz), High Gamma (65-170 Hz), Ultra-high Gamma (180-450 Hz), and the Multi-unit Activity Range (500-1,100 Hz).

**Figure 1:**
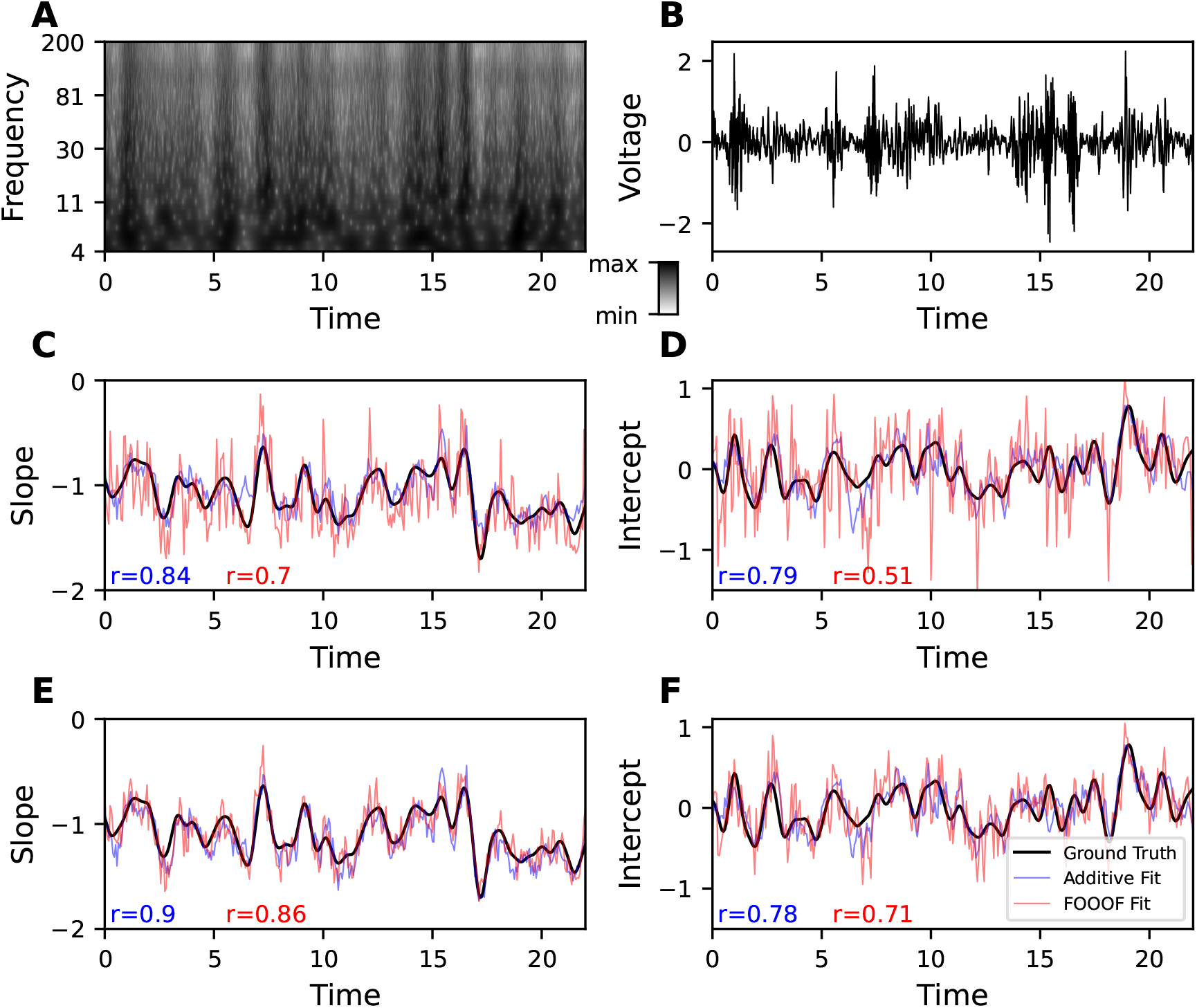
Additive regression recovers ground truth power-law parameters. **A**: Spectrogram representation of synthetic neural data as a function of time and frequency. **B**: Synthetic neural timeseries generated from spectrograms containing power-law and high-gamma components. **C**: Ground truth (black) and estimated power-law (blue for Additive and red for FOOOF, colors shared across panels) slopes as a function of time from the power-law plus high-gamma dataset. **D**: Ground truth and estimated power-law intercepts as a function of time from the power-law and high-gamma dataset. **E**: Ground truth and estimated power-law slopes as a function of time from the power-law only dataset. **F**: Ground truth and estimated power-law intercepts as a function of time from the power-law only dataset.

**Figure 2:**
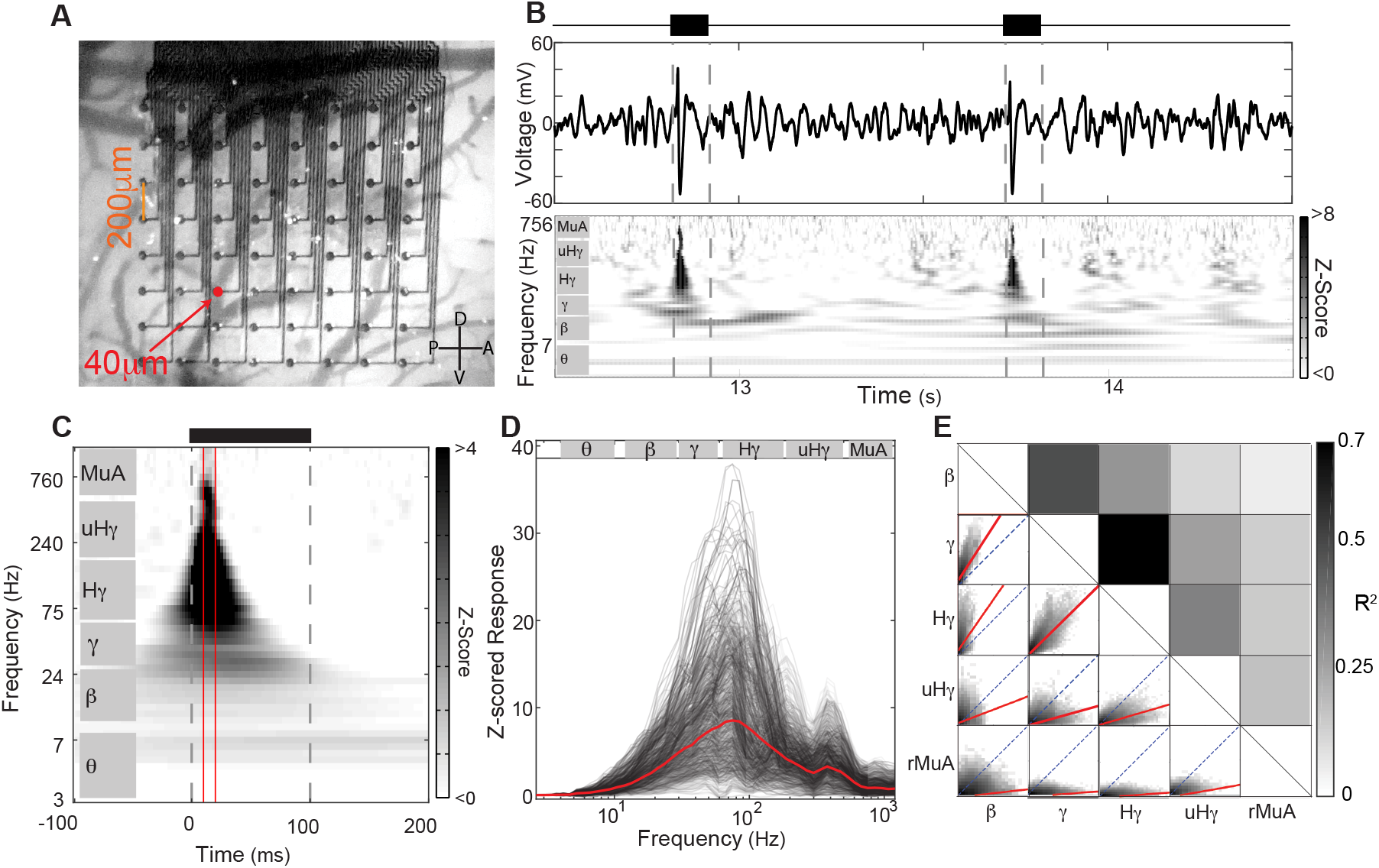
Stimulus evoked cortical surface electrical potentials contain multiple, distinct high-frequency peaks. **A**: Photomicrograph of an 8×8 *µ*ECoG grid (pitch: 200 *µ*m, contact diameter: 40 *µ*m) on the surface of rat primary auditory cortex (A1). **B**: Single-trial evoked cortical surface electrical potentials (top) and z-scored spectral decomposition (bottom) evoked by playback of white noise bursts extracted from a *µ*ECoG electrode over the rat primary auditory cortex. Stimulus onset and duration represented by black bar and grey dashed lines. Canonical banding of neural frequencies is indicated. **C**: Average neural spectrogram. Stimulus onset and duration represented by black bar and dashed lines. Canonical banding of neural frequencies is indicated. Red vertical lines in **C** corresponds to time-window of extracted evoked response used for subsequent analysis. **D**: Z-scored evoked response to white noise as a function of frequency for all active electrodes (active electrodes defined by peak z-score *>* 3.0; *N* = 1402). Black: individual electrodes, red: grand average across all electrodes. **E**: To quantify the relationship between different frequency bands, we performed pair-wise linear regressions between single-trial peak response magnitudes for different components of the evoked response (regressions were ordered as lower frequency predicting higher frequency). The results of this analysis are summarized with the median best-fit lines and coefficients of determination (*R*^2^) across all active electrodes (*N* = 1402). The slopes of the regressions (lower triangle, each cell displays a 2D histogram of data densities, red lines are median best-fit from linear regression on individual electrodes, blue line is unity), were ≥ 1 for lower frequencies predicting higher frequencies up to *Hγ*, and then reversed, with slopes *<* 1 for lower frequencies predicting higher frequencies greater than *Hγ*.

### Multi-unit triggered, average neural spectrograms

To more sharply focus analysis on the high-frequency CSEP components, we calculated multi-unit triggered, average neural spectrograms (MuTANS) by averaging the z-scored (see above) neural spectrogram centered (‘triggered’) at the time of each detected multi-unit event (**Fig. 3**), similarly to Logothetis et al. 2013. We applied a standard multi-unit thresholding procedure to form multi-unit events (Quiroga et al. 2004). In line with traditional usage from intra-cortical recordings (e.g., Lin et al., 2015), we use the term ‘thresholded multiunit activity’ (tMuA) for un-sorted, large-amplitude threshold crossings of fast (i.e., spike-like) voltage events, and term the corresponding waveforms multi-unit events to reflect the fact that they do not originate from a single neuron. For subsequent analysis of MuTANS (see below), we ensured non-negativity by adding the absolute value of the minimum of each MuTANS to that MuTANS (recall that MuTANS are constructed from z-scored neural spectrograms).

**Figure 3:**
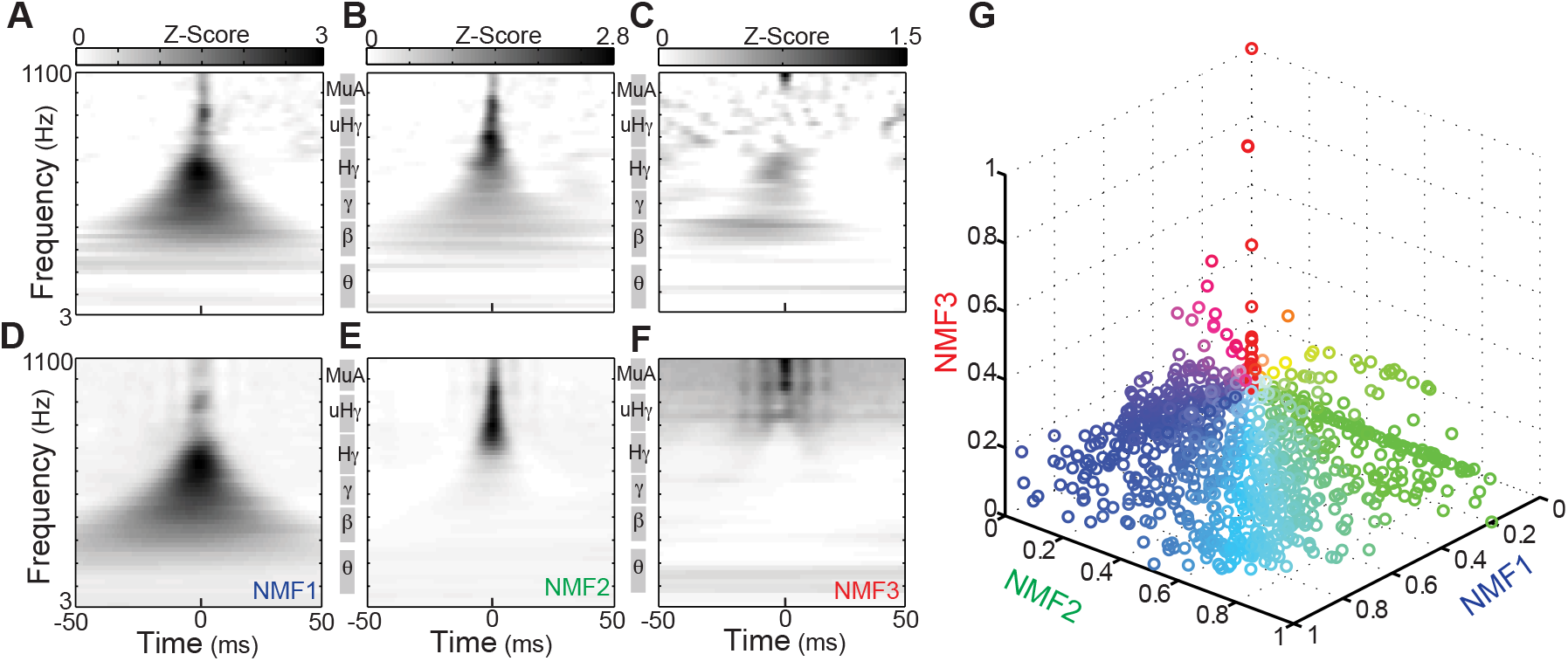
Cortical surface electrical potentials are composed of distinct high-frequency components. **A-C:** Example Multi-unit Triggered, Average Neural Spectrograms (MuTANS) (z-scored power at frequency as a function of time from multi-unit event detection) at three electrodes. We observed a diversity of MuTANS, with some having broadly distributed power focused in the *γ*-*Hγ* bands (**A**), others having narrower band power centered in the u*Hγ* range (**B**), and still others with even more narrow band power occurring at the highest frequencies measured **C. D-F:** Three UoI-NMF_cluster_ bases with relatively narrow band distributions of energy that correspond well to the example MuTANS displayed in **A-C. G:** Scatter plot of the normalized weights associated with NMF1-3 in the reconstruction of each data point (open circles, *N* = 1156). Each data point is colored RGB [NMF3, NMF2, NMF1] in proportion to the relative magnitude of the weight assigned to that NMF basis. Note the existence of ‘pure’ MuTANS, with contributions almost entirely from one NMF basis (pure red, green, and blue circles), lots of data points with large relative NMF1 (dark blue), and many points that are mix of NMF1 and NMF2 (light blue).

### The UoI-NMF_cluster_ algorithm

We analyzed the structure of MuTANS using our recently introduced, noise-robust non-negative matrix factorization (NMF) algorithm (UoI-NMF_cluster_, Ubaru et al. 2017). While a detailed description of this algorithm is outside the scope of this manuscript, here we provide the motivation, outline the innovations, and summarize the main statistical result. In brief, NMF algorithms aim to decompose a non-negative data matrix 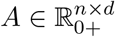 (ℝ_0+_ denotes the non-negative real numbers), into a basis matrix 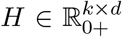 and a weight matrix 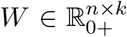, where *k* is the desired rank (i.e., number of bases):

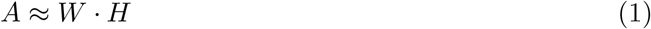

It has been observed that if the *n* data samples are generated by combining points on a simplex (i.e., the bases *H* are the vertices of a simplex), NMF algorithms will give a unique ‘parts-based’ decomposition of the data (see references in Ubaru et al. 2017). However, the parts-based decomposition of the basic NMF model (Eq 1) can be unstable if the data is corrupted by noise, as is often the case in ECoG data.

To overcome this shortcoming of basic NMF, we have recently introduced the UoI-NMF_cluster_ algorithm. This novel algorithm first learns a set of bases (*H*) across a set of bootstrap samples. The bases are clustered using dbSCAN, and the stability selection criterion is used to choose the “best” rank *k* (see references in Ubaru et al. 2017). Together, these innovations yield more accurate parts-based decompositions of noisy data and high-accuracy reconstructions of the de-noised data. See Ubaru et al. 2017 for further details on the algorithm and results on synthetic and other experimental data.

### Human speech data

The dynamics of human speech unfold at relatively fast timescales, in both the articulatory and neural domains. It is also known that the high-gamma amplitude is predictive of spoken speech at fast timescales. Human ECoG data recorded during speech production provides an oppourtunity to study the origin of the time-varying high-gamma signal in CSEPs. Here, we use a previously published dataset (Bouchard and Chang 2019). The experimental protocol, collection, and analysis of the human data have been previously described in detail (Bouchard and Chang 2014a; Bouchard and Chang 2014b; Bouchard, Mesgarani, et al. 2013; Livezey, Bouchard, et al. 2019). The experimental protocol was approved by the Human Research Protection Program at the University of California, San Francisco. The subjects gave their written informed consent before the day of surgery.

Four native English speaking human subjects underwent chronic implantation of a subdural electrocortigraphic (ECoG) array over the left hemisphere as part of their clinical treatment of epilepsy (**Fig. 4**). The subjects read and spoke-aloud consonant-vowel syllables (CVs) which were composed of 1 of 19 consonants followed by 1 of 3 vowels. Since subjects did not produce CVs with equal frequency, CVs with fewer than 10 repetitions were excluded per-subject. Across subjects, the number of repetitions per CV varied from 10 to 105, and the total number of usable trials per subject was S1: 2576, S2: 1584, S3: 4627, and S4: 1421.

**Figure 4:**
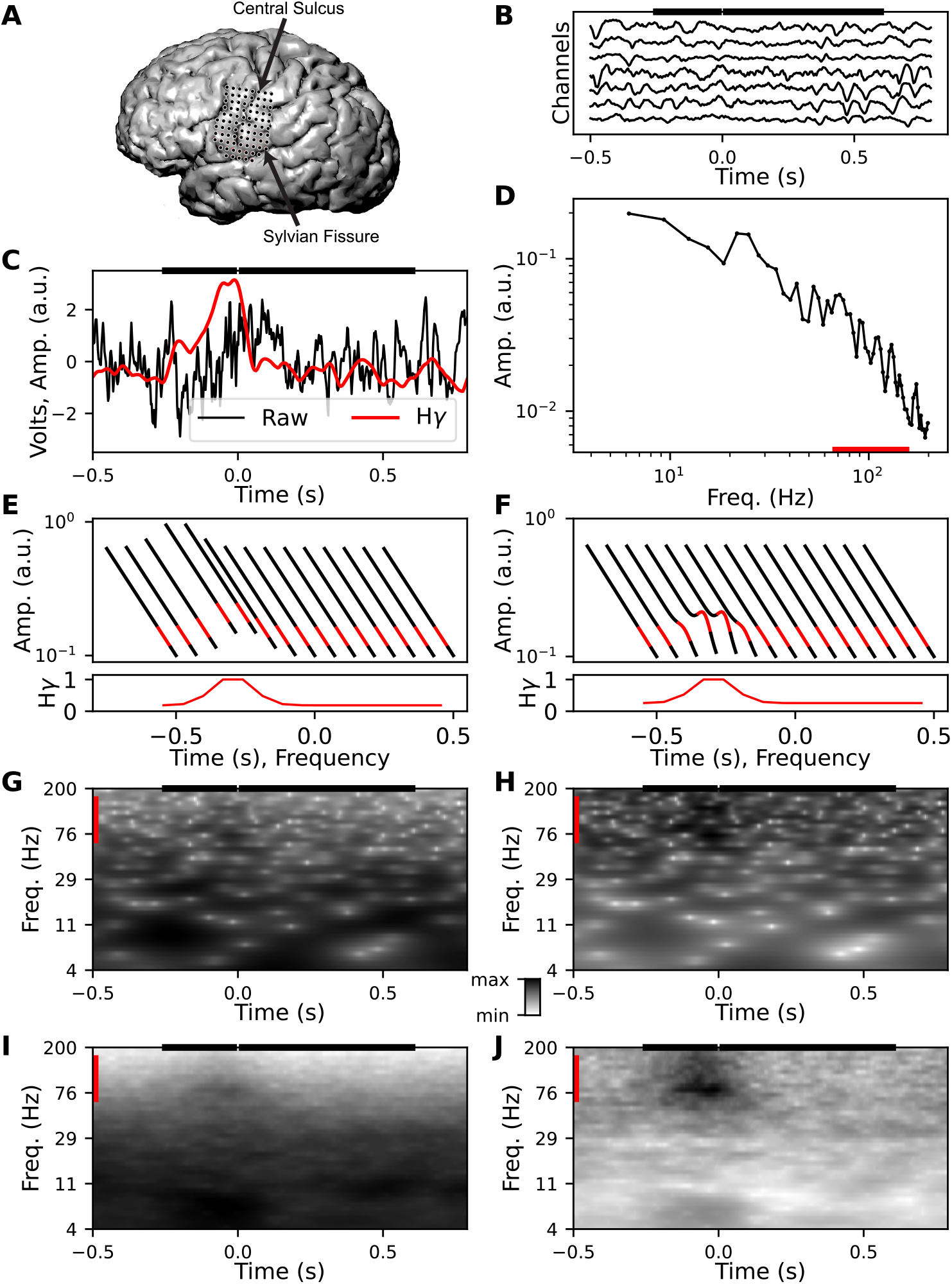
The time-varying high-gamma amplitude appears spectrally and temporally localized. High-gamma frequency range is shown in red along frequency axes. Approximate consonant and vowel timing is shown in black along the top of time axes. **A:** Anatomical reconstruction of Subject 1 with electrodes in the vSMC indicated. **B:** Timeseries for a subset of the electrodes during the production of “gaa”. **C:** Voltage timeseries on a single electrode for a single trial (black). The high-gamma amplitude is shown in red for the same trial and electrode. **D:** Average power spectrum for this window of data plotted on a log-log scale. **E, F:** Synthetic power spectra showing high-gamma modulation (bottom panels) arising from a broadband power-law component modulating in time (**E**) or a bandlimited high-gamma component modulating in time (**F**). Each line represents the power spectrum on a log-log scale at one point in time. The red section indicates the high-gamma range. **G:** Single-trial wavelet log-power as a function of frequency and time for the same electrode and trial as **C. H:** Single trial, baseline normalized wavelet log-power for the same electrode and trial. **I:** Trial averaged (*N* = 44) wavelet log-power as a function of frequency and time for the same electrode and CV. **J:** Trial averaged (*N* = 44), baseline normalized wavelet log-power for the same electrode and CV.

Cortical surface electrical potentials (CSEPs) were recorded directly from the cortical surface with a high-density (4mm pitch), 256-channel ECoG array and a multi-channel amplifier optically connected to a digital signal processor (Tucker-Davis Technologies, TDT, Alachua, FL). Channels with artifacts or excessive noise based on visual inspection were excluded from all subsequent analysis. The raw CSEP signals from the remaining channels were downsampled to 400 Hz in the frequency domain and then common-average referenced and 60 Hz line noise was removed. For each channel, the time-varying, constant-Q wavelet amplitude was extracted in the frequency domain at 40 logarithmically-increasing center frequencies between 4 and 200 Hz. Finally, the amplitudes were downsampled to 200 Hz. For analysis on the high-gamma band (70-150 Hz), individual bands with center frequencies in the range were z-scored to baseline and averaged to produce a single, normalized high-gamma amplitude. Based on previous results (Bouchard and Chang 2014a; Bouchard and Chang 2014b; Bouchard, Mesgarani, et al. 2013; Livezey, Bouchard, et al. 2019), we focused on the electrodes in the ventral sensorimotor cortex (vSMC). The number of usable electrodes in vSMC per subject was: S1: 86, S2: 78, S3: 83, and S4: 99. The activity for each of the examples in our data set was aligned to the acoustic onset of the consonant-to-vowel transition. For each example, a window 0.5 seconds preceding and 0.79 seconds following the acoustic onset of the consonant-to-vowel transition was extracted.

### Robust regression methods

To estimate fast timescale, power-law components in the human ECoG data, we required a robust estimation method that could account for the noisy, fast timescale, single trial variability. Linear regression can be used to fit power-laws to log-transformed data (**Fig. 5**). In this model, the log-amplitude (or equivalently log-power, up to a linear re-scaling) at each frequency is modeled as a linear function of the log-frequencies

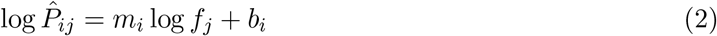

where *i* indexes the different trials or time points for a specific electrode and *j* indexes the frequencies.

**Figure 5:**
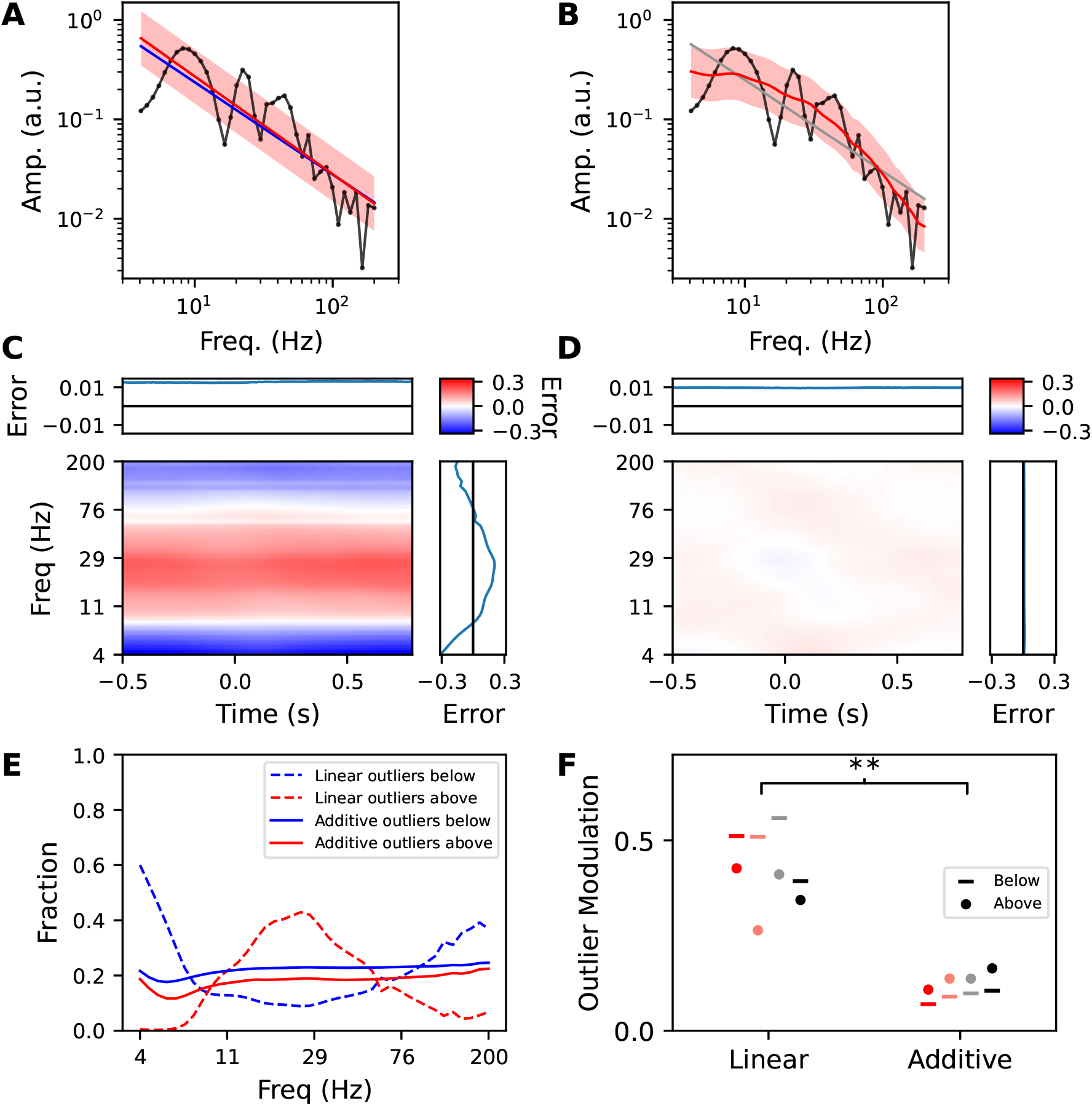
Additive robust regression removes bias from power-law estimates. **A**: Single trial wavelet power at one point in time (black line and dots). OLS regression fit in blue. Huber fit in red with shaded outlier boundary. **B**: Same neural data as **A**. Gray line is the linear component of the Additive Huber model. Red line is the linear and additive components with shaded outlier boundary. **C and D**: Each group of plots shows the median model error as a function of time and frequency, time, or frequency. The medians are taken across the remaining variables including trials and electrodes. Color-scale and corresponding axis limits are shared across **C** and **D. C:** Median errors for the Linear model fit with the Huber objective. **D:** Median errors for the Additive model fit with the Huber objective. **E:** For the Linear and Additive models fit with the Huber loss, the fraction of points that are classified as outliers above and below the regression line are plotted as a function of frequency for Subject 1. **F:** Across subjects (colors) and models the differences between the maximum and minimum outlier fraction across frequencies are shown.

Robust regression has been suggested for estimating broadband components (Manning et al. 2009). In regression problems, robustness can be gained by using the Huber loss (rather than the typical mean-squared error, MSE)

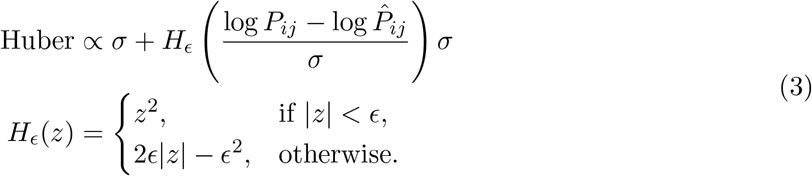

The MSE loss would place the same squared-error penalty on all points. This means that outliers, which have large errors, will have a large impact on the estimated regression coefficients. The Huber loss is more robust to outliers since in addition to fitting the regression coefficients, it also estimates a threshold *σ* at which errors switch from a MSE loss to a linear absolute value loss, which is less sensitive to outliers.

### Additive robust regression

Purely linear power-law regression cannot account for shoulders that are common in neural spectra. This limitation means that robust linear methods will conflate outliers due to noise or bandlimited components with points that deviate from the linear fit due to the shoulder. The Additive model includes an extra additive term for each electrode and frequency, *c*_*j*_, which is shared across time and trials and can model static shoulders in the spectrum (**Fig. 5**)

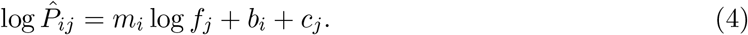

Compared to the number of parameters from *m*_*i*_ and *b*_*i*_, which are unique to every electrode, time, and trial, the inclusion of *c*_*j*_ represents a very small increase in the total number of model parameters. Each electrode gets 40 additional parameters compared to 2 parameters × 258 time points × approximately 2,000 trials per subject, which is about 10^6^ total parameters from *m*_*i*_ and *b*_*i*_. However, the additional structured flexibility greatly increases the model’s ability to fit shoulders in the neural spectra. This formulation of the Additive model is overparameterized. To resolve this, we choose the solution with the minimum *L*_2_ norm of *c*. This is a desirable choice because it forces *m* and *b* to account for as much of the structure in the neural spectra as possible. All models were created in Pytorch and optimized using L-BFGS-B from SciPy (Paszke et al. 2019; Virtanen et al. 2020).

We validated the Additive model in synthetic data. Power-law spectrograms were generated, as in Eq 8, in the time-frequency domain (**Fig. 1A**) then converted into timeseries data (**Fig. 1B**) through an inverse short-time Fourier transform. The ground truth, time-varying slopes (*m*_*i*_), intercepts (*b*_*i*_), and log-high-gamma amplitudes were drawn from Gaussian processes with squared-exponential kernel with timescales, means, and variances set to be similar to what we observed in the human ECoG data (**Fig. 1C-D**, black lines, high-gamma amplitude not shown). Data from a pure power-law plus high-gamma component (**Fig. 1C, D**) and power-law only (**Fig. 1E, F**) was considered. The spectrogram was given random uniform phase, and the shoulder (*c*_*j*_) was fixed as a function of time with shape estimated from data. Finally, a small amount of white noise was added (SNR = 1000).

We validated the Additive model by fitting the time-varying slope and intercept parameters. For comparison, we also estimate the slope and intercept parameters using FOOOF (Donoghue et al. 2020) although the Additive model and FOOOF require different preprocessing pipelines (wavelet amplitudes with log-spaced frequencies versus spectrograms with linear-spaced frequencies). We find that the Additive model recovers the power-law slope with high pearson correlation for the synthetic dataset with power-law and high-gamma components (**Fig. 1C**, Additive: *r* = 0.84, FOOOF: *ρ* = 0.70) and for the power-law only dataset (**Fig. 1E**, Additive: *ρ* = 0.90, FOOOF: *ρ* = 0.86). For the intercept, the choice of windowing functions in the spectrogram and wavelet amplitude calculations leads to an global scaling (additive offset in log-amplitude) in the total power which means it is difficult to recover the mean of the intercept although the variations in time can be recovered. Due to this, we mean center all ground truth and estimated intercepts for plotting, and the pearson correlation is insensitive to the mean. We find that the Additive model recovers the power-law intercept (up to a global additive offset) with high pearson correlation for the synthetic dataset with power-law and high-gamma components (**Fig. 1D**, Additive: *ρ* = 0.79, FOOOF: *ρ* = 0.51) and for the power-law only dataset (**Fig. 1F**, Additive: *ρ* = 0.78, FOOOF: *ρ* = 0.71). These results show that the Additive model can be used to recover power-law slope and intercept parameters.

There are no hyperparameters in the Linear or Additive robust regression models, so cross validation is not needed for hyperparameter selection. However, overfitting can be assessed and controlled by cross validation. In order to assess the errors and outliers in the Linear and Additive power-law models (**Fig. 5**), we performed a 10 fold set of fits to control for overfitting. In each fit, 10% (4 out of 40) of amplitudes across 40 frequencies are held-out as a test set, that is, they are masked during training. This masking is done independently for each channel, time, and trial. We then calculated the errors and counted outliers separately for the train and test sets and report the test set errors and outliers. In analysis that uses the Additive model parameters to reconstruct power-law spectra, we average *m, b*, and *c* across the 10 folds (**Figs. 6 and 7**).

**Figure 6:**
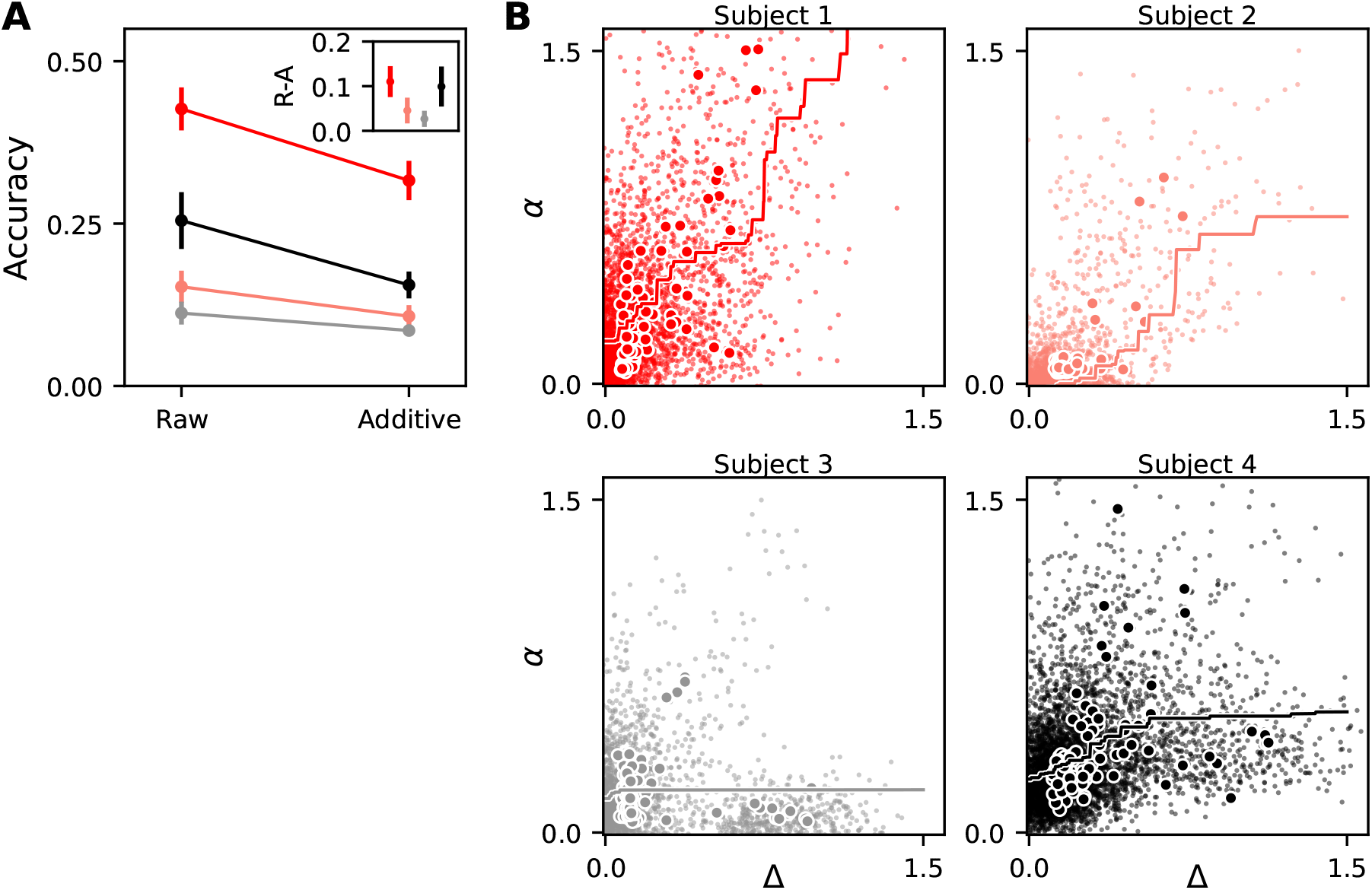
Classification of produced speech is driven by bandlimited high-gamma on a small number of electrodes. **A:** The test-set accuracy (mean ± std. across 10 cross validation folds) of a logistic regression classifier is shown for raw high-gamma amplitude (left) and high-gamma amplitude estimates from the Additive fits (right). The main panel shows the absolute classification accuracies and the inset panel shows the raw minus Additive (R-A) accuracy, compared across subjects and folds. Color is Subject ID. (same as in **B**). **B:** The error of the estimated log high-gamma amplitude (Δ) is scattered against the classification importance (*α*) for each Subject. Small dots are individual electrode-CV pairs and large circles are electrodes. Trendline is estimated using isotonic regression. Δ and *α* have been 0-1 normalized at 0% and 95% averaged across subjects to make the plotting scale less sensitive to a small number of outlier electrode-CVs outside of the plotting window.

**Figure 7:**
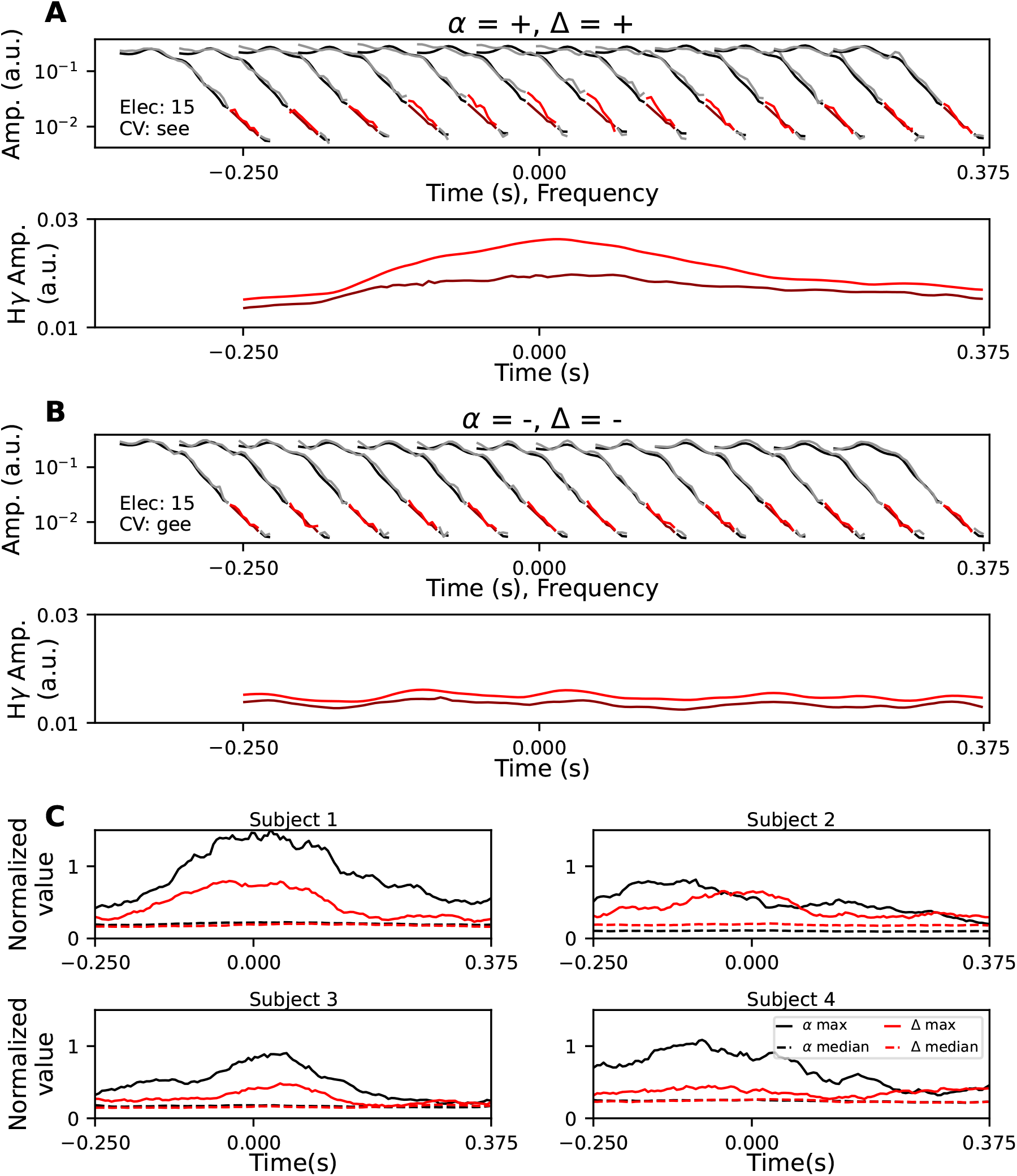
A bandlimited high-gamma component increases power-law fit errors during informative times. **A:** The trial-averaged amplitude across frequencies at a decimated set of times, windowed around the consonant-vowel acoustic transition (250 ms before to 375 ms after the transition) is shown in the top panel (maroon and black are raw data and grey and red are Additive fits). Time is aligned to the high-gamma portion of the spectrum. The average high-gamma amplitude (maroon and red) is shown in the bottom panel. The electrode shown was selected based on maximizing the variability in fits and importance across CVs. The CV with maximum Δ and *α* is shown. **B:** For the same electrode in **A**, the CV with minimum Δ and *α* is shown. **C:** The median of the top 1% of electrode-CVs (max) ranked by classificaiton importance and median across all electrode-CVs (median) is plotted for *α* and Δ across electrodes and CVs are shown as a function of time for each subject. *α* and Δ were 0-1 normalized to their minimum 0% and maximum 97.5% across time jointly for all subjects.

This type of mask-based cross validation is often required in unsupervised methods where some or all of the model parameters are fit per-sample, such as the scores matrix in principal components analysis or tensor components analysis (Williams et al. 2018). Entire trials or time points cannot be used as a test set since each trial and time point has its own *m* and *b* parameters. Likewise, a frequency cannot be used as a test set since *c* is shared across all frequencies in the Additive model.

### Logistic classification and error comparison

Determining the relationship between consonant-vowel (CV) prediction importance and goodness-of-fit in the high-gamma range as a function of time, ECoG electrode, and CV class gives insight into whether the high-gamma amplitude is solely generated from a broadband, power-law component. To determine CV predictive importance, we trained regularized logistic regression models to predict the CV class from single-trial, baseline-normalized high-gamma amplitudes and calculate the normalized model weights (*β*). To determine the goodness-of-fit in the high-gamma range, we look at the trial-averaged errors from the power-law fits in the high-gamma range (**Figs 6, 7**).

For classification, the dataset was split by single trials into 80-10-10 training, validation and testing sets across 10-folds with non-overlapping test sets. The validation accuracy was used to select the optimal regularization strength and the test set accuracy is reported. The features used in the regression were normalized as follows. We have additive power-law fits, 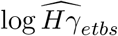, for the dataset, log *Hγ*_*etbs*_, where *e* indexes electrode, *t* indexes time, *b* indicates the recording block, and *s* indexes the sample or trial. In general, averages could be taken across time, channels, etc. We report the RMS of the trial-averaged, log normalized *Hγ* error from the power-law fits, Δ, as

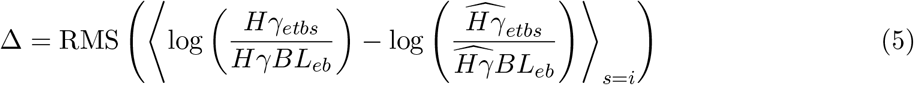

where *i* is the CV-label and *b* represents the recording block (each block has its own baseline, log *HγBL*_*eb*_, and baseline fit, log 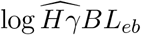 for every electrode). The RMS can be taken over one or more dimensions (CV, time, electrodes) and plotted as a function of the remaining dimensions.

We fit logistic regression models with *L*_2_ (ridge) regularization on single trial, baseline-normalized amplitudes, 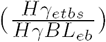. Typically, regression features are z-scored to the training statistics if interpretable *β* values are desired. However, normalizing the raw amplitudes by their mean baseline value makes the amplitudes more interpretable (removes arbitrary scaling due to electrode contact or positioning) and typically increases the classification accuracy of regularized logistic regression models. However it means that the weights of the logistic regression model (*β*) are difficult to interpret since the features used in the regression were not z-scored. To combine the benefits of baseline normalization and z-score normalization, we trained the models with baseline normalized features and then re-scaled the *β* values using the standard deviations, making the final *β* values interpretable. This 2-stage normalization impacts how *L*_2_ regularization changes features with differing amplitudes normalized to baseline. Electrodes with large amplitude relative to baseline will have larger final *β* values with the 2-stage scheme compared to if they had been z-scored before the regularized regression. We report the regression importance, *α*, as the RMS of the regression weights for the logistic regression model trained on the raw high-gamma amplitudes

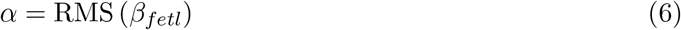

where *f* indexes cross-validation folds, *e* indexes electrode, *t* indexes time, and *l* is the CV-label. The RMS can be taken over one or more dimensions (folds, CV, time, electrodes) and plotted as a function of the remaining dimensions.

The values for both Δ and *α* can be compared across subjects and electrodes. Since Δ is normalized to baseline, the power-law fit errors have a meaningful scale across both electrode and subjets. For *α*, the variance of the neural inputs is taken into account after performing regression; however, the regression targets (CV labels) cannot be z-scored or normalized (unlike in linear regression), and there is some variation in both the total number of CV classes and the distribution of trials per CV class across subjects. Although the variability is not large, this may reduce the absolute comparability for *α* across subjects, although this does not impact within-subject electrode comparisons.

### PCA analysis

Miller, Zanos, et al. 2009 calculate the ensemble normalized log-power 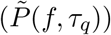 across frequencies, trials, and times, using their notation

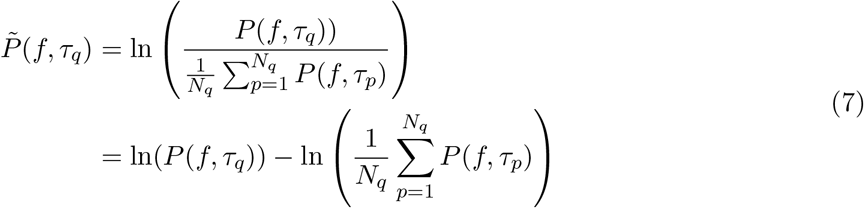

where *f* indexes the frequencies and *τ*_*q*_ indexes the trial and time (**Fig. 8**). They then use the log-powers as the features and trials and times as the samples. Note that although the “ensemble normalization” standardizes the features (divides the raw powers by their mean), the average is inside of the log in the second term, which does not lead to mean subtractions of the log-powers.

**Figure 8:**
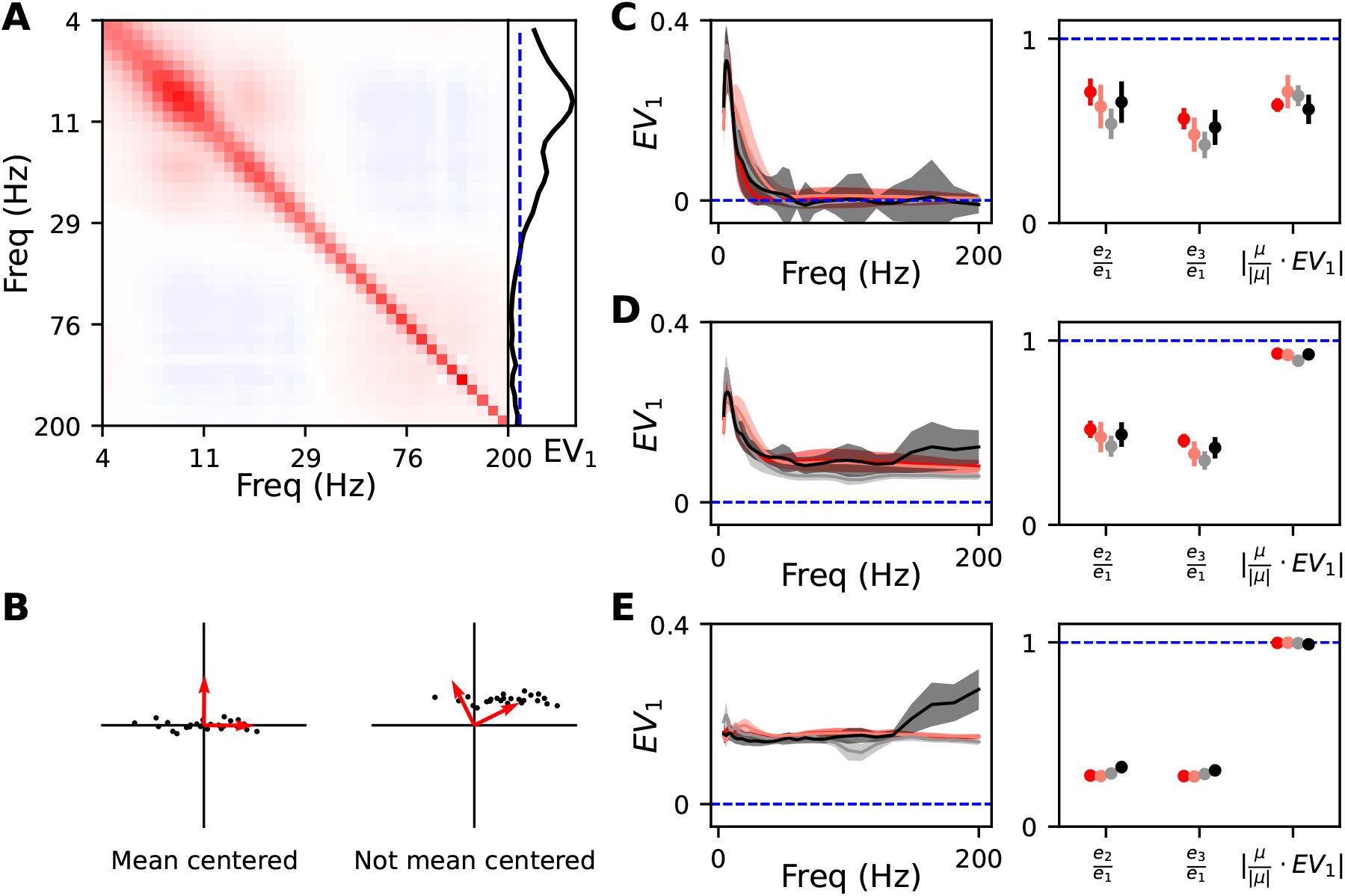
The highest variance spectral component is bandlimited and low frequency. **A:** Cross-frequency covariance matrix for an electrode from S1 (red is positive correlation, blue is negative). The elements of the first eigenvector are shown to the right (blue dashed line is zero). **B:** The black dots are drawn from a 2d synthetic dataset with axis-aligned principal components. The red arrows are the eigenvalues derived from the covariance matrix on the left and the second moments matrix on the right. **C-E** For each pair of panels (left and right), the left panel shows the median across electrodes of the first eigenvector’s elements and the 25th and 75th percentiles for all subjects (colors indicate Subject ID). The right panel shows the ratio between the first and second eigenvalues, the ratio between the first and third eigenvalues, and the absolute cosine angle between the first eigenvector and the unit-normalized mean. **C:** Eigen analysis applied to the log-power data (mean-centered) covariance matrix. **D:** Eigen analysis applied to the log-power data (not mean-centered) second moment matrix. **E:** Eigen analysis applied to the trial-shuffled log-power data (not mean-centered) second moment matrix.

### Data Availability

The rat data used in this study will be made available upon publication. The human data used in this study is available at https://doi.org/10.6084/m9.figshare.c.4617263.v4 (Bouchard and Chang 2019).

### Code and Software Accessibility

UoI-NMF_cluster_ is available as part of the PyUoI library (Sachdeva et al. 2019). The Additive model and related analysis will be made available upon publication.

## Results

Cortical surface electrical potentials (CSEPs) are composed of many broadband and band-limited sources (Buzsáki, Anastassiou, et al. 2012). The high-gamma range (70-150 Hz) is traditionally thought to be a single broadband component whose power is driven by spiking activity (Manning et al. 2009; Miller 2010; Miller, Sorensen, et al. 2009). However, the timescales over which neural spectra are often averaged and the noise floor of clinical amplifiers has made it difficult to probe the structure of time-varying, high frequency (70 - 1000 Hz) CSEPs in many studies. Therefore, the structure of high frequency CSEPs at fast timescales is poorly understood. Here, recorded evoked CSEPs in rates and humans and developed a novel robust regression method to characterize the high frequency structure of CSEPs recorded in rats and in humans. In rats, we analyzed the structure of CSEPs evoked by white noise bursts using *µ*ECoG and showed that the high frequency range is componsed of several distict high frequency components. In humans, we analyzed CSEPs recorded during a speech production task which allowed us to show that the variability in the high-gamma range cannot be accounted for by a purely broadband component, and that the broadband component has less predictive power than the raw high-gamma amplitude. Finally, we show that the highest variance component in CSEP spectra is low-frequency and bandlimited and show that improper data normalization led previous studies to misinterpret their results as a broadband component.

### Stimulus evoked cortical surface electrical potentials contain multiple, distinct high-frequency peaks

In order to characterize the high frequency range of CSEPs, we recorded large amplitude, fast-timescale CSEPs from auditory cortex in response to the presentation of bursts of auditory white noise in rats (see Materials and Methods). An example of a 64-channel grid is shown on the auditory cortical surface (subdural placement) of an anesthetized rat in the photomicrograph in **Fig. 2A**. An example recording from the electrode demarcated in red in **Fig. 2A** during two consecutive stimulus presentations is shown in **Fig. 2B** (stimulus onset and offset indicated by grey dashed vertical lines). The top panel displays the wide-band cortical surface electrical potential (10 - 1100 Hz), while the bottom panel displays the corresponding normalized neural spectrogram. For each recording electrode and spectrogram frequency, we z-scored the time-varying amplitude relative to baseline (see Materials and Methods for details). **Fig. 2C** shows the average normalized evoked neural spectrogram derived from the recorded electrical potential in response to this stimulus (*N* = 60 trials). For both single trials, as well as the average, white noise stimuli evoked large-amplitude, rapid CSEP deflections. Indeed, responses in the gamma (*γ*: 30-60 Hz) and high-gamma (*Hγ*: 70-150 Hz) ranges were very large, exceeding four standard deviations of the baseline activity in this example. Strikingly, we also observed large responses in the higher frequency bands that extended up to 1100 Hz.

To summarize the frequency content of evoked CSEPs across all recordings, we extracted mean z-scored responses for all spectrogram frequencies in a ±5ms window around the peak response time (defined by the median of peak response times for each frequency, red vertical area in **Fig. 2C**). We defined a responsive electrode as a having peak z-score across frequencies *>* 3.0 (*N* = 1402*/*1664 electrodes (84%) from 17 *µ*ECoG placements on auditory cortex in 6 rats). **Fig. 2D** plots the z-scored response as a function of frequency for all responsive electrodes (black: individual electrodes, red: grand average across all electrodes). We found a highly non-uniform response profile across frequencies, with extremely large magnitude responses in the *γ/Hγ* range (∼40 - 150 Hz; z-score *>* 20 in many cases), as well as a pronounced secondary peak in the ultra high-gamma (u*Hγ*) range (200 - 450 Hz), and robust (z-score *>* 3.5) responses in the multi-unit activity (MuA) range (*>* 500*Hz*) at many recording sites. These results confirm the single-trial robustness of the peaked nature of the response as a function of frequency around *γ*/*Hγ* observed in **Fig. 2D**. Across all electrodes, the peak *R*^2^ between the mean evoked neural spectrogram and single-trial spectrograms (of the same duration) during the stimulus presentation had a median of 0.650 [0.323 0.808, 25/75 CI]. In contrast, the peak *R*^2^ between the mean evoked neural spectrogram and single-trial spectrograms (of the same duration) during the non-stimulus times had a median of 0.034 [0.008 0.096; 25/75 CI]. These distributions were significantly different: *p <* 10^*−*20^, *N* = 1402, WSRT. On single-trials, we observed only a modest relationship between the relative magnitudes of *Hγ* and MuA at the same electrode (**Fig. 2E**, *<* 20% variance shared, *R*^2^ = 0.18 from linear regression). Indeed, the amount of shared single-trial variance across frequency components at the same electrode declined with increasing difference in frequency (**Fig. 2E**, *R*^2^ = 0.75, slope = −2.4 (log units), *P <* 0.005, *N* = 10). This raises the possibility that high-frequency evoked CSEPs can be decomposed into distinct spectral components.

### Cortical surface electrical potentials are composed of distinct high-frequency components

We next examined the degree to which high-frequency content of CSEPs can be decomposed into distinct components. For each channel, we calculated multi-unit triggered average neural spectrograms (MuTANS) by averaging the neural spectrogram centered at the time of each detected multi-unit event (see Materials and Methods). The plots in **Fig. 3A-C** display example MuTANS (z-scored power at frequency as a function of time from event detection) at three electrodes. We observed a diversity of MuTANS, with some having broadly distributed power focused in the *γ/Hγ* bands (**Fig. 3A**), others having narrow-band power centered in the u*Hγ* range (**Fig. 3B**), and still others with even more narrow band power occurring at the highest frequencies measured (MuA, **Fig. 3C**). To summarize the diversity of MuTANS, we used our recently introduced, noise-robust non-negative matrix factorization algorithm UoI-NMF_cluster_ (Ubaru et al. 2017) (see Materials and Methods for details) to decompose the entire collection of measured MuTANS into a smaller number of “generator” MuTANS that can be additively combined to reconstruct the observed data (see Materials and Methods). **Fig. 3D-F** display three bases (of seven, sorted by explained variance) learned by UoI-NMF_cluster_ with distributions of power that correspond well to the example MuTANS displayed in **Fig. 3A-C** (NMF 1 : *γ/Hγ*; NMF 2 : u*Hγ*; NMF 3 : MuA), indicating that these profiles are robustly present across the entire data set.

To quantitatively determine the degree to which these three NMF bases were present in the experimental data, we max-normalized the weights associated with NMF bases 1-3 in the reconstruction of each data point (*N* = 1156). This normalization removes variations in the magnitudes (i.e., z-scores) of MuTANS across electrodes, emphasizing the relative degree to which the three NMFs are present in the MuTANS. The scatter plot in **Fig. 3G** displays the normalized weights associated with NMF bases 1-3, where each data point is colored blue-green-red in proportion to the relative magnitude of the weight assigned to NMF bases 1-3, respectively. We observed many MuTANS were primarily composed of NMF 1 (dark blue) or were an approximately equal mix of NMF1 and NMF2 (light blue). Of particular note, we found many “pure” MuTANS which were composed almost entirely of either NMF1, NMF2, or NMF3 (pure dark blue, green, and red circles along the axes). These results demonstrate that high-frequency evoked cortical surface electrical potentials can consist of distinct spectrotemporal components (NMF1 : *γ/Hγ*; NMF2 : u*Hγ*; NMF3 : MuA), and thus are not broadband phenomena.

### The time-varying high-gamma amplitude appears spectrally and temporally localized

During speech production, both articulatory and neural dynamics unfold over relatively fast (∼ 100 ms) timescales. The high-gamma power at these timescales has been shown to be highly predictive in a number of speech (produced and perceived) decoding tasks (Anumanchipalli et al. 2019; Bouchard and Chang 2014b; Livezey, Bouchard, et al. 2019; Livezey and Glaser 2021; Mugler et al. 2014; Yang et al. 2015), with task engagement typically localized in time and on a subset of electrodes (Bouchard, Mesgarani, et al. 2013). Cortical surface electrical potentials (CSEPs) have been observed to approximately have power-law scaling as a function of frequency (Manning et al. 2009; Miller 2010; Miller, Sorensen, et al. 2009) well into the high-gamma (*Hγ*) range. In the *Hγ* band, it is currently not clear whether the raw power should be considered to be broadband (power-law) component plus noise (Miller, Zanos, et al. 2009) or a combination of a broadband component, a band-limited component, and noise at fast timescales. CSEPs recorded during the production of speech allow us to test whether the time-varying high-gamma amplitude only contains a broadband component, as opposed to a combination of broadband and bandlimited components.

Cortical surface electrical potentials were recorded from the ventral sensorimotor cortex (vSMC) of four human subjects (**Fig. 4A** shows vSMC electrodes from Subject 1, B shows an example single trial recording from several electrodes). The raw timeseries (**Fig. 4C**, black line) was converted to a spectrotemporal representation with a 40 band wavelet transform, including the high-gamma range (**Fig. 4C**, red line). Single trial CSEP amplitudes display a power-law trend as a function of frequency (**Fig. 4D**). **Figure 4E** shows illustrated data where the a time-varying broadband component (top panel in E, each line is a log-log plot of the neural amplitude as a function of frequency, time and frequency increase from left to right) is the sole component in the time-varying high-gamma amplitude (red lines, average high-gamma amplitude in the bottom panel of E). This is in contrast to an alternative scenario in **Figure 4F**, where an illustrated time-varying bandlimited component in the high-gamma range (top panel in F) is combined with a broadband component in the time-varying high-gamma amplitude. Without the larger frequency context of the power-law fits, it is not possible to distinguish the source of variations in high-gamma amplitude (**Fig. 4E, F** bottom panels).

The power-law trends and single-trial variability obscure the variation in the high-gamma range (**Fig. 4G**), although per-frequency baseline normalization (**Fig. 4H**), trial averaging (**Fig. 4I**), and their combination (**Fig. 4J**) reveals clear high-gamma modulations (note that baseline normalization removes the power-law structure). This indicates that there may be a robust, bandlimited high-gamma component that modulates independently from variability in the power-law component in the high-gamma range.

### Additive robust regression removes bias from power-law estimates

Linear regression between the log-power and log-frequency has been used to estimate power-law broadband components in neural spectra (Manning et al. 2009; Miller, Sorensen, et al. 2009). However, since CSEP spectra are believed to be a combination of broadband and band-limited components, simple OLS regression will be potentially biased by band-limited components. In addition, additive Gaussian noise in the original time-domain signal can lead to heavy tails towards negative amplitude in the distribution of log-amplitudes (light spots in **Fig. 4G**) (Rivet et al. 2007). To be less sensitive to these potential outliers and tails, robust regression has been previously used to fit the underlying power-law component (Manning et al. 2009; Owen 2007). Using the Huber loss to fit a power-law component also gives an estimated error threshold to switch between a MSE loss (*L*_2_) and a robust absolute value loss (*L*_1_) for “outliers” (**Fig. 5A**, black dots and lines are raw data, red line is the estimated power-law component, red shaded area is the outlier threshold, blue is is the OLS estimated power-law).

Using linear Huber regression, we find that there are large systematic changes in the distribution of median model errors across frequency (**Fig. 5C**, center and right panels), although the distribution of median model errors from the fits is stationary across time (**Fig. 5C** top panel). Due to the non-stationarity across frequency, the fraction of data points estimated to be outliers in the Huber loss have large modulation across frequencies (**Fig. 5E** dashed lines, outliers are grouped by whether then are above or below the threshold). This modulation in median error and outliers means that the model is not consistently classifying deviant points in the spectrum as outliers as a function of frequency, which will cause bias in the fit power-law parameters.

We developed a new power-law estimation model that has the robust properties of Huber regression with an additional additive term which mitigates the bias across frequency. The curvature observed in the spectra comes from a combination of one or more shoulders in the spectrum and the wavelet filter widths. Let 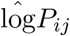 be the estimated log-power for a particular electrode at each wavelet center log-frequency log *f*_*j*_ (indexed by *j*) across trials and time (both indexed by *i*). To account for the curvature, we can fit a linear model with a frequency specific additive offset, *c*_*j*_

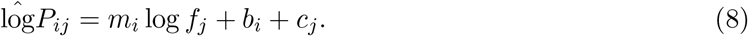

Here, each time, trial, and electrode has its own *m* and *b* estimate that is shared across frequencies. Each frequency and electrode has its own *c* estimate that is shared across time and trials. The *c* parameter additively “couples” regressions that were previously independent in the purely linear model. This leads to a power-law regression model that has a (time-varying) linear component (**Fig. 5B**, grey line), but also can fit a static shoulder which influences both the final fit parameters and outliers thresholds (**Fig. 5B**, red line and red shaded area, respectively). Materials and Methods has the details of the model description, fitting procedure, and validation on synthetic data.

When fit to neural spectra, the Additive robust regression model has relatively small modulation of the median regression error as a function of frequency compared to the linear model (**Fig. 5D**, same colormap and scale as C). The fraction of points classified as outliers has significantly smaller modulation across frequency for the additive model (**Fig. 5E**, solid lines). Across subjects and outliers above and below the threshold, the modulation across frequency (measured by the max minus min values from **Fig. 5E**) is significantly smaller for the Additive model compared to the Linear model as (**Fig. 5F**, two-sided, Wilcoxon signed-rank test, T=36, p=0.0078, rank-biserial correlation=1.0). This shows that, in addition to inheriting the robust properties of linear Huber regression, the Additive Robust regression model can account for shoulders, a common feature of neural spectra, enabling better estimation of the underlying power-law component.

### Classification of produced speech is driven by bandlimited high-gamma on a small number of electrodes

We hypothesized that if the high-gamma (*Hγ*) amplitude is only made up of a power-law component and noise, denoising the raw amplitudes using the power-law fits should give a better estimate of the high-gamma amplitude. Using the denoised high-gamma amplitude may increase the consonant-vowel (CV) classification accuracy and should not decrease the classification accuracy depending on the original signal-to-noise ratio. The functional organization of vSMC leads to sparse activity during the production of consonant-vowel syllables (Bouchard, Mesgarani, et al. 2013). This means that we should expect the inferred electrode importances to be highly heterogeneous across time and syllables when predicting produced speech. We expect the electrodes with high functional engagement with the task (as determined through classification importance) to provide the clearest signal as to whether the informative high-gamma component can be explained by a power-law fit.

To test this hypothesis, we trained logistic regression models to classify the spoken syllables from the high-gamma amplitudes. We compared the classification accuracy on the test set when the models were trained on the raw high-gamma amplitudes (**Fig. 6A**, left points, color indicates subject) versus the high-gamma amplitudes from the Additive fits (**Fig. 6A**, right points, see Materials and Methods for details). We find that across subjects and cross validation folds, classification accuracy is significantly higher on the raw high-gamma amplitudes than on the high-gamma amplitude from the Additive fits (**Fig. 6A**, inset shows the differences in accuracies, two-sided Wilcoxon signed-rank test, T=820, p=3.6×10^*−*8^, rank-biserial correlation=1.0). This shows that the raw high-gamma amplitude is more informative than the high-gamma amplitude from the Additive power-law fit.

In order to assess whether the most highly informative electrodes have the best (or worst) fits from the Additive robust regression model, we calculated the classification importance (*α*) for the logistic regression model trained on the raw high-gamma amplitudes and the high-gamma fit error (Δ) with RMS calculated across cross validation folds and time for *α* and time for Δ (**Fig. 6B**, small dots). The RMS was additionally calculated across CVs, giving *α* and Δ per electrode (**Fig. 6B**, large dots). Larger *α* indicates higher classification importance and larger Δ indicates larger error in the power-law fits. Qualitatively, most electrode-CVs have both low classification importance and low high-gamma fit error. However, electrode-CVs and electrodes that have high classification importance tend to also have larger high-gamma fir error. For all subjects, we found positive Spearman correlations between *α* and Δ calculated across electrodes and CVs (Spearman correlations are S1: 0.46, S2: 0.17, S3: 0.10, S4: 0.38. All p-values *<* 10^*−*10^). This shows that electrode-CVs that are more important for classification also tend to have higher high-gamma fit error. Subjects 1 and 4 have more electrodes and electrode-CVs that have high classification importance relative to Subjects 2 and 4 which is consistent with Subjects 1 and 4 having higher classification accuracy. Subjects 3 and 4 both have a group of electrodes (lower-right part of the panels) that have low classification importance and large fit errors. This may indicate that those electrodes have large amounts of noise. Finally, we observe that the order of Spearman correlations is the same as the order of classification accuracies (S1, S4, S2, S3, from highest to lowest). Together, these results demonstrate that the Additive fits are not denoising a power-law high-gamma component that is solely predictive of spoken speech syllables, indicating that high-gamma amplitudes have a task informative, bandlimited high-gamma component.

### A bandlimited high-gamma component increases power-law fit errors during informative times

Due to the temporal structure of the consonant-vowel (CV) production task, it is expected that the high-gamma amplitudes are most informative around *t* = 0, that is, around the acoustic consonant-vowel transition (Bouchard, Mesgarani, et al. 2013; Livezey, Bouchard, et al. 2019). If the high-gamma amplitude is only a combination of power-law amplitude and noise, points in time where the evoked high-gamma amplitude is informative should increase the signal-to-noise ratio, leading to better high-gamma fits (lower Δ), at least for electrodes that are informative for the task. On the other hand, if the informative high-gamma component is from a bandlimited high gamma component, potentially in addition to the power-law component, informative times should have worse high-gamma fits (higher Δ) as they will deviate more from the power-law component.

To qualitatively understand the temporal relationship between classification importance and high-gamma fit error, we examined the raw data and fits as a function of time and frequency. We selected the electrode from Subject 1 that had the highest variability of classification importance (*α*) and highest variability of high-gamma fit error (Δ) across CVs (Electrode 15). We then chose the CVs with the highest *α* and Δ (**Fig. 7A**) and lowest *α* and Δ (**Fig. 7B**). Electrode 15 is very important for classifying the CV “see”. Before or after *t* = 0, the Additive fits closely match the raw data (**Fig. 7A**, top panel, gray and red lines are the raw trial-averaged data, black and maroon lines are the Additive fits, frequency increase from left to right within a spectrum, time increase from left to right across spectra). However, the timecourse of high-gamma amplitude and the power-law fit in the high-gamma range shows a high error peaked around *t* = 0 (**Fig. 7A**, averages across frequencies in the high-gamma range is shown in the bottom panel). Conversely, the same electrode is not important for predicting the CV “gee” although the high-gamma fits are very good. The Additive fits closely match the raw data across time and frequency (**Fig. 7B**, top panel), including in the high-gamma range (**Fig. 7B**, bottom panel). This highlights the heterogeneity of classification importance and high-gamma fit errors.

To quantitatively understand the relationship between classification importance and high-gamma fit error as a function of time, we can look at the time-course around *t* = 0 (−0.25 seconds to 0.375 seconds) of *α* and Δ. We summarize *α* and Δ for the 1% maximally important electrode-CVs (“*α* max” and “Δ max”) and all electrode-CVs (“*α* median” and “Δ median”, respectively) by taking medians of either the 1% or all electrode-CVs respectively. We find that as *α* max peaks near *t* = 0, *α* median stays relatively constant (**Fig. 7C**, solid black and dashed black lines, respectively). This confirms the expected result that task engagement is highly sparse across electrode-CVs. However, the high-gamma fit error for the same “max” electrode-CVs (Δ max) is also at or considerably above the median (Δ median) across all electrode-CVs (**Fig. 7C**, solid red and dashed red lines, respectively). Furthermore, the timecourse of *α* max and Δ max are significantly positively correlated during this time window across subjects (Spearman correlations, S1: 0.87, S2: 0.57, S3: 0.86, S4: 0.30, all p-values *<* 10^*−*3^). This indicates that the errors in Additive high-gamma fits are being driven by a highly informative, bandlimited component in the high-gamma range.

### The highest variance spectral component is bandlimited and low frequency

Broadband components will have the largest impact on the interpretation of bandlimited components if they contribute to a large fraction of the cross-frequency variance across time. In one extreme, broadband components could be the largest source of variance in neural spectra (Miller, Zanos, et al. 2009). In this case, the first principal component could be used to estimate the broadband component, rather than estimating the broadband component using (Additive) power-law fitting. In this section we investigate whether PCA can be used to estimate broadband components.

To determine whether broadband fluctuations exist in the data as the main source of variance, we performed a PCA analysis similar to Miller, Zanos, et al. (2009) (also used in Miller, Honey, et al. 2014) on the full cross-frequency covariance matrix (**Fig. 8A**, left panel, red is positive covariance, blue is negative). We compute the log of the ensemble-normalized power across frequencies (*f*) as features for PCA 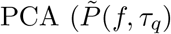, see Materials and Methods for details) for each electrode, using trials and time (*τ*) as samples. For each subject and electrode, we compute the first principal components (PC_1_ or equivalently eigenvector of the covariance EV_1_ shown in **Fig. 8A**, right panel black line) and flip its sign such that its largest element is positive. We also compute the first 3 eigenvalues. The ratios of the first-to-second and first-to-third eigenvalues are used to determine whether EV_1_ is well separated from the other eigenvectors, indicating that it is the largest single source of variance rather than part of a larger subspace with nearly uniform variance.

Unlike Miller, Zanos, et al. (2009), we mean-center the features before forming the covariance matrix and performing the eigen-decomposition, as is done in PCA. We believe that leaving the mean in the data likely led to a misinterpretation of the meaning of EV_1_. Miller, Zanos, et al. (2009) compute an eigen-decomposition on the product of 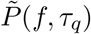 and its transpose (i.e., the second moments matrix)

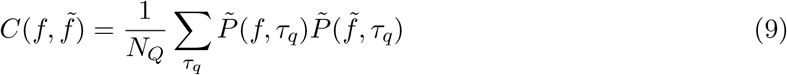

rather than on the covariance matrix

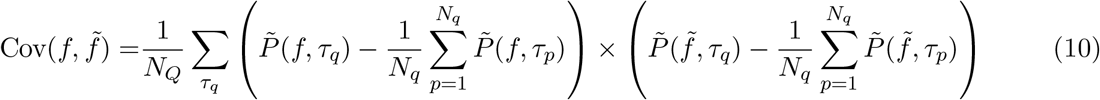

where the feature means have been subtracted. Since 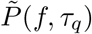 will generally have nonzero mean, using Eq 9 versus Eq 10 will generally give different eigenvectors which require different interpretations. In comparing these two matrices, it is useful to distinguish PC_1_, the first principal component of the covariance matrix and EV_1_, the first eigenvector of any matrix.

To gain intuition for the difference in synthetic data, we show that not mean subtracting the data biases EV_1_ away from PC_1_. Samples from 2-dimensional multivariate normal data generated with axis-aligned eigenvalues (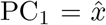 with eigenvalue 1 and PC_2_ = *ŷ* with eigenvalue .025) are shown in **Fig. 8B**. If we mean-center the data, we recover the true principal components which are axis-aligned (**Fig. 8B**, left panel). However, if we perform an eigen-decomposition on the second moments matrix (Eq 9), we find that PC_1_ is corrupted by the mean data vector and is biases towards the center of the point cloud rather than pointing along the x-axis (**Fig. 8B**, right panel). This makes EV_1_ uninterpretable as a source of variability because it mixes static features (the mean) with features that are varying (the true PCs) and for this reason it cannot be used to estimate time-varying broadband components.

Across subjects and electrodes, we do not find consistent broadband structure in the first principal components (EV_1_). The elements of the first eigenvector (EV_1_) are typically largest for frequencies less than 25 Hz and are close to zero for larger frequencies (**Fig. 8C**, left panel, colors indicate subjects as in **Fig. 6**). We do not find roughly constant nonzero EV_1_ elements as was reported in Miller, Zanos, et al. 2009 (compare Figure 1A from (Miller, Zanos, et al. 2009) to **Fig. 8C**) that could be interpreted as broadband variation. The first eigenvalue is also well seperated from the second and third, confirming that it is not parted of a larger subspace of similar variance (**Fig. 8C**, right panel). The mean and EV_1_ are correlated, but not colinear (**Fig. 8C**, right panel). These results show that the highest variance spectral component is bandlimited and low frequency.

When we do not mean-center our data before performing an eigen-decomposition, we indeed find a component in EV_1_ (**Fig. 8D**, left panel) where all elements of the EV_1_s are non-zero and have the same sign. However, the EV_1_s we find using this method are almost perfectly collinear with the mean of the data (**Fig. 8D**, right panel). The impact of not mean-centering can be exaggerated by trial-shuffling the features. In this case, trial-shuffling destroys any correlation across features and so PC_1_ should be aligned to the single feature with the largest variance. However, when the shuffled data is not mean centered, EV_1_ still retains a “broadband component” even though all cross-frequency correlations have been removed (**Fig. 8E**, left panel) and EV_1_ is almost perfectly correlated with the mean (**Fig. 8E**, right panel). Together, these analyses show that broadband components are not the main source of time-varying variance in the neural spectra.

These results show how previous studies likely misinterpreted their results by using the second moments matrix rather than the covariance matrix in the eigen-decomposition. However, this analysis does not exclude the existence of broadband components that may still be present in our data and mixed into (multiple) lower magnitude principal components, although it is unlikely that PCA will be useful for extracting broadband components if this is the case.

## Discussion

In this work, we tested whether the high-frequency range (70-1,000 Hz) of cortical surface electrical potentials (CSEPs) are composed of multiple broadband and bandlimited components, and in particular, whether the the high-gamma range (70-150 Hz) contains a time-varying bandlimited component. We recorded evoked CSEPs with *µ*ECoG in rats and used a previously published dataset of ECoG recordings in humans. In rats, we showed that there are multiple evoked peaks in power in the gamma/high-gamma, ultra high-gamma, the multi-unit activity range. Furthermore, we showed that multi-unit triggered average neural spectrograms, can be decomposed into components that have bandlimited, temporally localized power in the high-gamma, ultra high-gamma, the multi-unit activity range. In humans, we showed that the raw high-gamma amplitude was significantly more predictive of speech than the power-law component in the high-gamma range. This indicates that there is a bandlimited component in the high-gamma range. Together, these results demonstrate that there are multiple bandlimited components in the high-gamma and high frequency range of CSEPs.

The high-gamma range of high frequency CSEPs is of particular interest because it is a highly informative signal for clinical applications of brain-computer interfaces, and it is related to multi-unit spiking activity (Baratham et al. 2022; Livezey and Glaser 2021; Ray and Maunsell 2011). In CSEPs recorded from rat auditory cortex, we observed multiple peaks in the evoked high frequency neural spectra. Across single-trials, the correlations between these peaks decreased as the difference in their frequency increased, indicating that they were not generated by the same biophysical source. Indeed, a robust non-negative matrix factorization algorithm decomposed the multi-unit triggered average neural spectrograms into components that have temporally sparse, bandlimited structure. Previous work has shown that the high-gamma amplitude can be used to classify spoken consonant-vowel (CV) syllables with high accuracy (Bouchard and Chang 2014b; Bouchard, Mesgarani, et al. 2013; Livezey, Bouchard, et al. 2019). Here, we leveraged this to discern whether time-varying, power-law fits to CSEPs would denoise an underlying power-law component in the high-gamma range and lead to higher classification accuracy. Contrary to this, we found that the high-gamma power-law fits had lower classification accuracy compared to the raw high-gamma amplitude, indicating that the power-law fits discarded a predictive bandlimited signal. We also showed that a small number of electrodes have high classification importance, which peaked near the consonant-vowel acoustic transition, and that these electrodes tend to have larger high-gamma fit errors. This is consistent with a bandlimited high-gamma component that causes both sparse functional importance and poor power-law fits. These results support the hypothesis that time-varying high-gamma amplitude arises from at least two distinct components: a power-law component and a bandlimited component. More broadly, we conjecture that the generating sources of the distinct frequency components observed in rat CSEPs may be spiking activity in different layers of cortex (Baratham et al. 2022). Because neurons in different cortical layers perform different computations, if true, this conjecture would imply that different frequency components are biomarkers of layer-resolved cortical computations.

In rats and humans, we found that the amplitude of evoked CSEPs in the gamma and high-gamma ranges were highly correlated (Livezey, Bouchard, et al. 2019). This is despite the fact that stimulus evoked, transient high-gamma amplitude arises from synchronous spiking of neurons (likely pyramidal cells) in layers 5 and 6 (Baratham et al. 2022), while the gamma oscillation is tied to perisomatic inhibition and irregular spiking (Buzsáki and Wang 2012). Indeed, our prior results indicate that the stimulus evoked neural response that drives the high-gamma amplitude has frequency “spill-over” into the range associated with gamma band (Baratham et al. 2022). Thus, we believe it is important to distinguish from such ongoing, more sustained oscillations from the transient increases in power associated with evoked responses.

Developing models of time-varying neural spectra is an area of ongoing research (Donoghue et al. 2020; Manning et al. 2009; Miller 2010; Miller, Sorensen, et al. 2009). The Additive robust regression model proposed here was developed under the requirement that it fit power-law components robustly to single-trial, fast-timescale noisy data, be scalable to millions of fits without any by-hand tuning, and account for shoulders in the neural spectra. Although methods such as FOOOF (Donoghue et al. 2020) have been used to fit neural spectra taken over much longer time windows (which can have higher signal-to-noise ratios due to averaging), to our knowledge, no method exists that fits all of the requirements at hand. The Additive model combines the robust Huber loss with a set of additive parameters that can account for shoulders in the spectrum. We showed that the Additive model was able to recover ground-truth power-law parameters in synthetic data and had a more flat distribution of outliers across frequency in neural data.

It had previously been proposed that broadband variability was the highest variance principle component in neural spectra (Miller, Zanos, et al. 2009). However, as we recapitulated in the human data, leaving the mean of the features in while forming the second moments matrix (rather than the covariance matrix, which relies on mean-centered features) implies that the first eigenvalue and eigenvector of the matrix was likely corrupted by the static mean. When the covariance is used, we find that the highest variance spectral component is low-frequency and bandlimited. We expect future work looking to decompose spectra into bandlimited and broadband components to be fruitful although stronger assumptions (than highest variance) about the form or properties of the broadband component will likely be needed.

Future work could take advantage of several properties of spectral decompositions to potentially improve the estimates of the power-law components in neural data. First, spectrotemporal decompositions generally induce correlations in the estimates of power across frequency and time. These could be incorporated into a generative model of time-varying power-law spectra to improve inference (Turner and Sahani 2014). Furthermore, rather than relying on robust generalizations of the MSE (Gaussian noise model) for fitting, the distribution of amplitudes could be considered more directly and parameterized in terms of Rayleigh or Rice distributions (or their log-variants) (Siddiqui 1964; Talukdar and Lawing 1991) which could better models the asymmetries and tails of the distributions of (log) amplitudes. Rather than considering bandlimited signals as outliers to be ignored during the estimation of the power-law component, as we did here, the power-law and bandlimited signals could be jointly estimated in addition to a potentially time-varying shoulder as in (Donoghue et al. 2020). However, incorporating all of these features into a single statistical model would require further development of appropriate priors for the broadband and bandlimited features and development of appropriate joint inference methods.

Estimating components from neural spectra (e.g., power-laws or bandlimited oscilliations) is important to link the biophysical mechanismcs that produce CSEPs to the higher order cognitive processes they are used to understand. That is, fitting a power-law model or bandlimited peak to neural spectra does not directly solve the inverse problem of ascribing CSEP power to a particular biophysical source; however, it can provide more resolved biomarkers of neural activity with known relationships to biophysical processes (Baratham et al. 2022; Buzsáki, Anastassiou, et al. 2012; Buzsáki and Wang 2012; Donoghue et al. 2020; Miller, Sorensen, et al. 2009; Ray and Maunsell 2011). Likewise, many features in neural spectra have known relationships to cognitive processes (Agarwal et al. 2014; Bouchard, Mesgarani, et al. 2013; Engel and Fries 2010; Herrmann et al. 2004; Klimesch 2012). Improving the ability to accurately decompose time-varying neural spectra into more biophysically motivated, time-varying components will help bridge the gap between biophysical sources of voltage in the brain and processes like neurological disorders, attention, and behavior. Such methods could be used to test the hypothesis that the time-varying high frequency range of CSEPs has resolvable contributions from multiple cortical layers, which would give non-penetrating recording technologies such as ECoG access to neural signals across depth in cortical columns.

## Acknowledgements

JAL was supported by the LBNL LDRD “Deep Learning for Science” and the Weill Institute for Neuroscience at UCSF (Bouchard). AH was supported by the LBNL LDRD “Deep Learning for Science”. MED was supported by the LBNL LDRD “Neural Systems and Data Science Lab”. EFC was supported by NIDCD (5R01DC012379-10). KEB was supported by LBNL LDRD “Neural Systems and Data Science Lab” and NINDS R01(RNS118648A). We thank the Neural Systems and Data Science lab for helpful discussions.

